# High-Resolution Mapping of the Human E3-Substrate Interactome using Ubicon Uncovers Network Architecture and Cancer Vulnerabilities

**DOI:** 10.1101/2025.04.09.647947

**Authors:** Tomoya Sakuma, Yuki Otani, Hideyuki Shimizu

**Affiliations:** Department of AI Systems Medicine, M&D Data Science Center, Institute of Integrated Research, Institute of Science Tokyo, Tokyo, JAPAN; Graduate School of Medical and Dental Sciences, Institute of Science Tokyo, Tokyo, JAPAN

**Keywords:** E3 ubiquitin ligase, E3-substrate interaction (ESI), Multimodal learning, Network biology, Cancer genomics

## Abstract

Mapping the intricate network of E3 ubiquitin ligase-substrate interactions (ESIs), essential for cellular regulation and implicated in numerous diseases including cancer, remains a central challenge limiting mechanistic understanding. Existing experimental and computational methods suffer from limitations in throughput, accuracy, and contextual relevance. Here, we introduce Ubicon, a deep learning framework designed to overcome these hurdles for accurate, proteome-wide human ESI prediction. Ubicon achieves high fidelity by synergistically integrating multimodal features: sequence embeddings from a protein language model specifically adapted for ESI prediction via parameter-efficient fine-tuning (PEFT), predicted 3D structures, and subcellular localization context. Validated extensively, Ubicon achieves state-of-the-art performance (AUROC = 0.9305, AUPRC = 0.6812), significantly surpassing previous approaches. Applying Ubicon, we constructed a high-resolution map of the human E3-substrate interactome, revealing its systems-level architecture characterized by hub proteins and functional modules linked to distinct biological processes. Integrating predictions with cancer genomics further uncovered disease-specific ESI network rewiring, involving oncogenic ligases like AURKA and CDC20, linked to poor prognosis. Ubicon provides a powerful platform to dissect ubiquitin signaling, uncover disease mechanisms, and inform targeted protein degrader development.

## Introduction

The ubiquitin-proteasome system (UPS) functions as a cornerstone of eukaryotic cell biology, wielding precise control over protein quality and abundance to orchestrate a vast array of essential cellular processes, from cell cycle progression and signal transduction to DNA repair and immunity^1–3^. Its critical role is underscored by the numerous human diseases, including cancers, neurodegenerative disorders, and immune deficiencies, linked to UPS dysfunction^4–7^. At the heart of the UPS’s regulatory specificity lies the E3 ubiquitin ligase family, comprising over 600 distinct enzymes in humans, each tasked with recognizing a specific repertoire of substrate proteins^1,8^. These E3-substrate interactions (ESIs) weave a complex regulatory network—the "ubiquitin code"^9–11^—that dynamically shapes cellular behavior and fate. Deciphering this intricate network is therefore not only fundamental to understanding basic life principles and disease pathogenesis but also holds immense promise for identifying novel therapeutic vulnerabilities. Indeed, the rational design of powerful therapeutic modalities like Proteolysis-Targeting Chimeras (PROTACs), which co-opt specific E3 ligases to induce the degradation of pathological proteins, critically depends on detailed knowledge of ESI selectivity^12–14^. Despite this centrality and burgeoning therapeutic relevance, systematically mapping the proteome-wide ESI landscape remains a formidable bottleneck, hampered by the sheer scale of potential pairings (>600 E3s vs. >20,000 proteins) and the often transient, context-dependent nature of these molecular recognition events^1,15^.

Efforts to chart this landscape have traditionally relied on experimental methods like yeast two-hybrid (Y2H) and Affinity Purification-mass spectrometry (AP-MS)^16–19^. While foundational, these approaches are fundamentally constrained by limitations in throughput and sensitivity, often failing to capture low-affinity or transient interactions critical for physiological signaling^20^. Furthermore, their susceptibility to experimental artifacts (false positives/negatives) and the challenge of replicating native cellular environments can obscure the true dynamics of the ESI network *in vivo*^21–24^. Complementary computational strategies have emerged, yet many remain confined to recognizing known short linear motifs (e.g., degrons), extrapolating from known domain-domain interaction paradigms, or utilizing general sequence-derived physicochemical features^25–29^. Such methods, while valuable for hypothesis generation, inherently struggle to predict truly novel interactions, particularly those mediated by complex, holistic structural interfaces rather than short motifs, and lack robust mechanisms to integrate crucial cellular context, such as subcellular compartmentalization or cell-state specific protein expression. Consequently, a comprehensive and accurate understanding of the human ESI network remains elusive.

The scientific landscape is currently being reshaped by breakthroughs in artificial intelligence (AI), presenting an unprecedented opportunity to address this long-standing challenge^30–32^. Large-scale protein language models (PLMs), pre-trained on the vast universe of protein sequences, have proven adept at capturing the complex statistical patterns and implicit biochemical information embedded within sequence data^33^. Concurrently, the advent of high-accuracy structure prediction algorithms, most notably AlphaFold2^34^, provides atomic-resolution insights into the three-dimensional architecture of proteins at a proteome scale. These powerful AI tools offer the potential for a paradigm shift in ESI prediction, enabling the integration of deep sequence context with detailed structural features – precisely the information modalities needed to overcome the limitations of previous methods. However, effectively harnessing these advances necessitates a bespoke computational framework capable of synergistically integrating these rich, multimodal data streams (sequence, structure, cellular context) and specifically optimizing them for the intricate task of predicting E3-substrate recognition.

Addressing this critical need, we introduce Ubicon, a novel deep learning framework architected to decode the human ESI network with high fidelity and resolution (Fig. 1). Ubicon’s core innovation lies in its sophisticated, synergistic integration of multimodal biological data. It fuses features derived from (i) state-of-the-art PLM embeddings, specifically refined for ESI prediction using parameter-efficient fine-tuning (PEFT) via Low-Rank Adaptation (LoRA)^35^ to capture nuanced sequence determinants; (ii) predicted 3D structural information, providing crucial insights into protein shape and potential interaction surfaces; and (iii) predicted subcellular localization, embedding essential cellular context (Fig. 1a and 1b). This integrated approach empowers Ubicon to achieve state-of-the-art predictive accuracy. Importantly, Ubicon’s biological relevance is demonstrated through *in silico* perturbation analyses, which reveal its sensitivity to the removal of known functional domains, indicating that it has learned biologically meaningful structure-function relationships. Leveraging this predictive power, we constructed the most comprehensive high-resolution map of the human ESI network to date, revealing novel insights into its systems-level organization and functional modularity. Furthermore, integrating Ubicon’s predictions with extensive cancer genomic and transcriptomic data allowed us to identify recurrent patterns of ESI network rewiring in human malignancies and correlate them with clinical outcomes (Fig. 1c). Ubicon thus provides a transformative platform to accelerate proteome-wide ESI mapping, deepen our mechanistic understanding of ubiquitin signaling in health and disease, and ultimately guide the development of novel therapeutic strategies targeting the UPS, including the optimization of next-generation targeted protein degraders.

**Figure 1.**
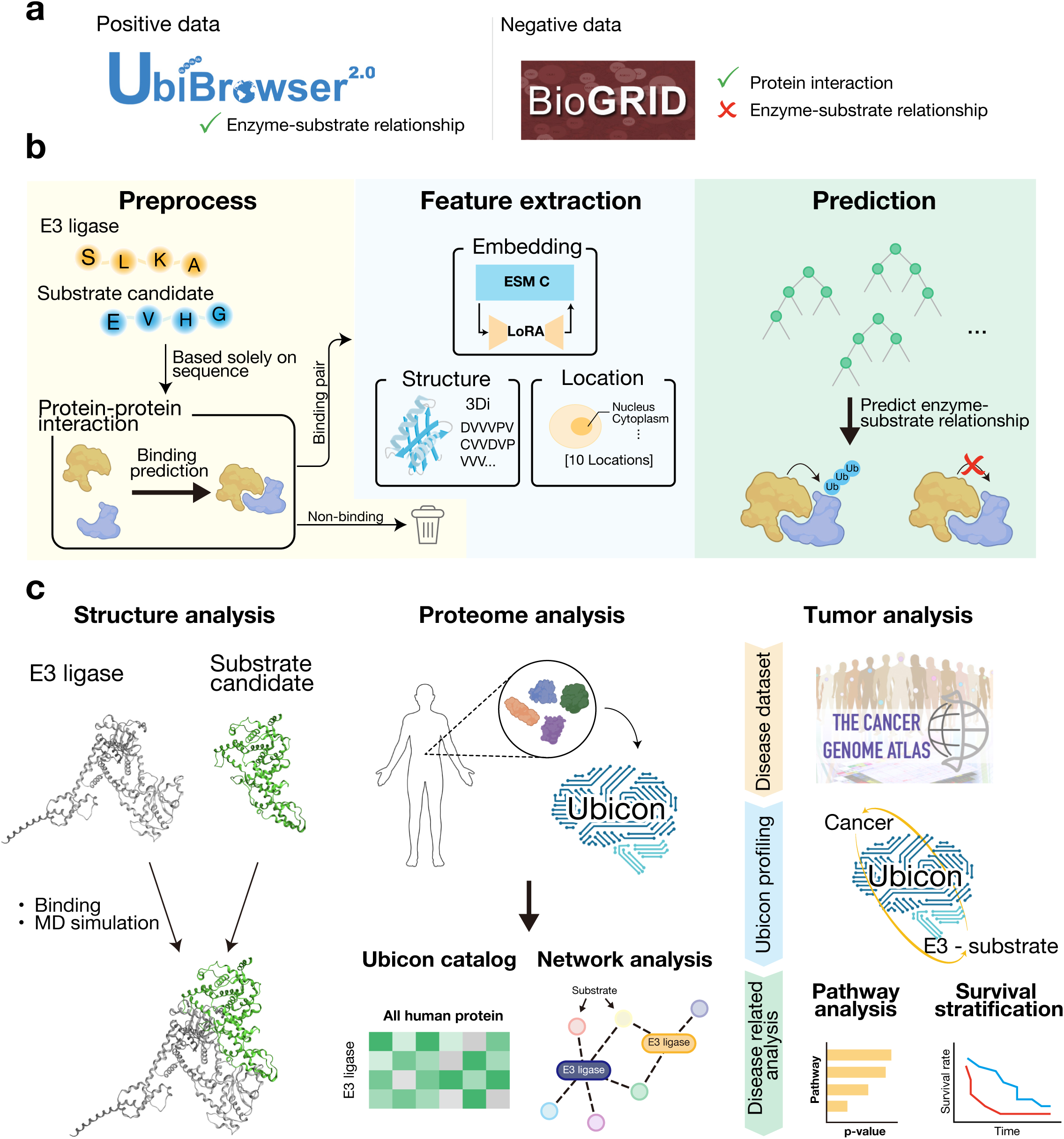
The Ubicon framework: Strategy for high-fidelity ESI prediction and systems-level biological discovery. (**a**) Curation of training datasets. The Ubicon model was trained using curated datasets. Positive examples, representing known E3-substrate interactions (ESIs), were obtained from the UbiBrowser 2.0 database. High-confidence negative examples were derived from known physical protein interactions documented in BioGRID, after carefully excluding known ESI pairs (see Methods for details). (**b**) Overview of the Ubicon prediction workflow. The Ubicon system predicts ESIs based solely on protein amino acid sequences as input. The workflow involves an initial Preprocessing step using protein-protein interaction (PPI) prediction to efficiently filter the vast search space to candidate pairs likely to physically interact. Subsequently, Feature extraction generates rich, multimodal representations for these candidate pairs, integrating deep sequence context from a task-adapted protein language model (ESM-C fine-tuned with LoRA), predicted structural information (3Di states), and predicted subcellular localization. Finally, these integrated features are fed into a machine learning classifier for high-accuracy prediction of ESI probability. Created with BioRender.com. (**c**) Applications and downstream analyses enabled by Ubicon. The high-fidelity ESI predictions generated by Ubicon serve as a powerful foundation for diverse downstream biological investigations explored in this study. These include: rigorous structural and functional validation to confirm that Ubicon captures known determinants of interaction (Structure analysis; see Fig. 3 for details); systems-level analysis of the proteome-wide ESI network to uncover its global architecture and organizational principles (Proteome analysis; see Fig. 4); and integration with disease datasets (e.g., TCGA) to investigate ESI network perturbations in human cancer and their clinical significance (Tumor analysis; see Fig. 5). Created with BioRender.com.

## Results

### Multimodal features and optimized architecture establish a high-performance ESI prediction baseline

Accurate prediction of E3-substrate interactions (ESIs) necessitates computational models that effectively integrate diverse biological information. Recognizing that ESIs depend on physical contact and three-dimensional structure, while many predictors rely primarily on sequence^25–29^, we first established an optimal feature set and model architecture.

We represented sequence information using embeddings from ESM-C^36^, a protein language model (PLM) capturing evolutionary context, and incorporated 3D structural information via 3Di state sequences derived from AlphaFold2^34^ structures using Foldseek^34,37^. To integrate these different modalities of information, we initially explored a CNN-based architecture (Fig. S1a). Comparing PLMs, the larger ESM2 (15B parameters) offered no significant performance increase over ESM-C (6B parameters) to justify its higher computational cost (Fig. S1b). Furthermore, ESM-C handles longer sequences, enabling broader applicability. We therefore selected ESM-C. Evaluating the contribution of structure, we found that adding 3Di structural information to ESM-C embeddings improved the Area Under the Precision-Recall Curve (AUPRC) (Fig. S1c). These results confirmed that combining deep sequence context (ESM-C) with structural information (3Di) provides a superior feature foundation for ESI prediction.

Building upon this optimized multimodal feature set (ESM-C 6B + 3Di), we benchmarked seven distinct machine learning architectures, using 5-fold cross-validation to identify the most effective ESI classifier. In addition to the initial CNN-based approach, we evaluated CatBoost^38^, XGBoost^39^, LightGBM^40^, Random Forests^41^, and Transformer-based tabular models (TabNet^42^, and TabPFN^43^). Among these, CatBoost achieved superior performance with the highest AUROC (0.9136), outperforming all other tested architectures (Table 1). This establishes ESM-C embeddings combined with 3Di structural features, classified by CatBoost, as a robust, high-performance baseline model for ESI prediction.

**Table 1:** Benchmarking machine learning models for ESI prediction with ESM-C 6B + 3Di features.

| ESI classifier | Mean AUROC ± SD | Mean AUPRC ± SD | Mean F1 Score ± SD |
| --- | --- | --- | --- |
| Initial CNN-based model | 0.8262 ± 0.0080 | 0.3143 ± 0.0316 | 0.3405 ± 0.0381 |
| CatBoost | 0.9136 ± 0.0040 | 0.5901 ± 0.0185 | 0.4480 ± 0.0212 |
| LightGBM | 0.9002 ± 0.0050 | 0.5803 ± 0.0173 | 0.4563 ± 0.0129 |
| XGBoost | 0.9050 ± 0.0048 | 0.5773 ± 0.0198 | 0.4880 ± 0.0219 |
| Random Forest | 0.9022 ± 0.0042 | 0.5037 ± 0.0154 | 0.4708 ± 0.0225 |
| TabNet | 0.7793 ± 0.0090 | 0.2712 ± 0.0214 | 0.3163 ± 0.0133 |
| TabPFN | 0.8422 ± 0.0060 | 0.3359 ± 0.0114 | 0.3753 ± 0.0080 |

### Boosting ESI prediction with adapted embeddings culminates in the high-accuracy Ubicon model

To further enhance predictive power, we hypothesized that fine-tuning PLM embeddings specifically for ESI recognition would outperform general pre-trained features. We employed parameter-efficient fine-tuning (PEFT) using Low-Rank Adaptation (LoRA)^35^ on the ESM-C 300M model, adapting its representations through contrastive learning on known ESI pairs (Fig. 2a, 2b). Remarkably, this LoRA-tuned ESM-C 300M model substantially outperformed the much larger, generally pre-trained ESM-C 6B model across all metrics (AUROC, AUPRC, F1 score; Fig. 2c). This result demonstrates that task-specific adaptation via LoRA unlocks critical features for ESI recognition within PLMs, achieving accuracy superior to even larger, generalist models and highlighting the importance of tailoring embeddings for complex molecular recognition tasks.

**Figure 2.**
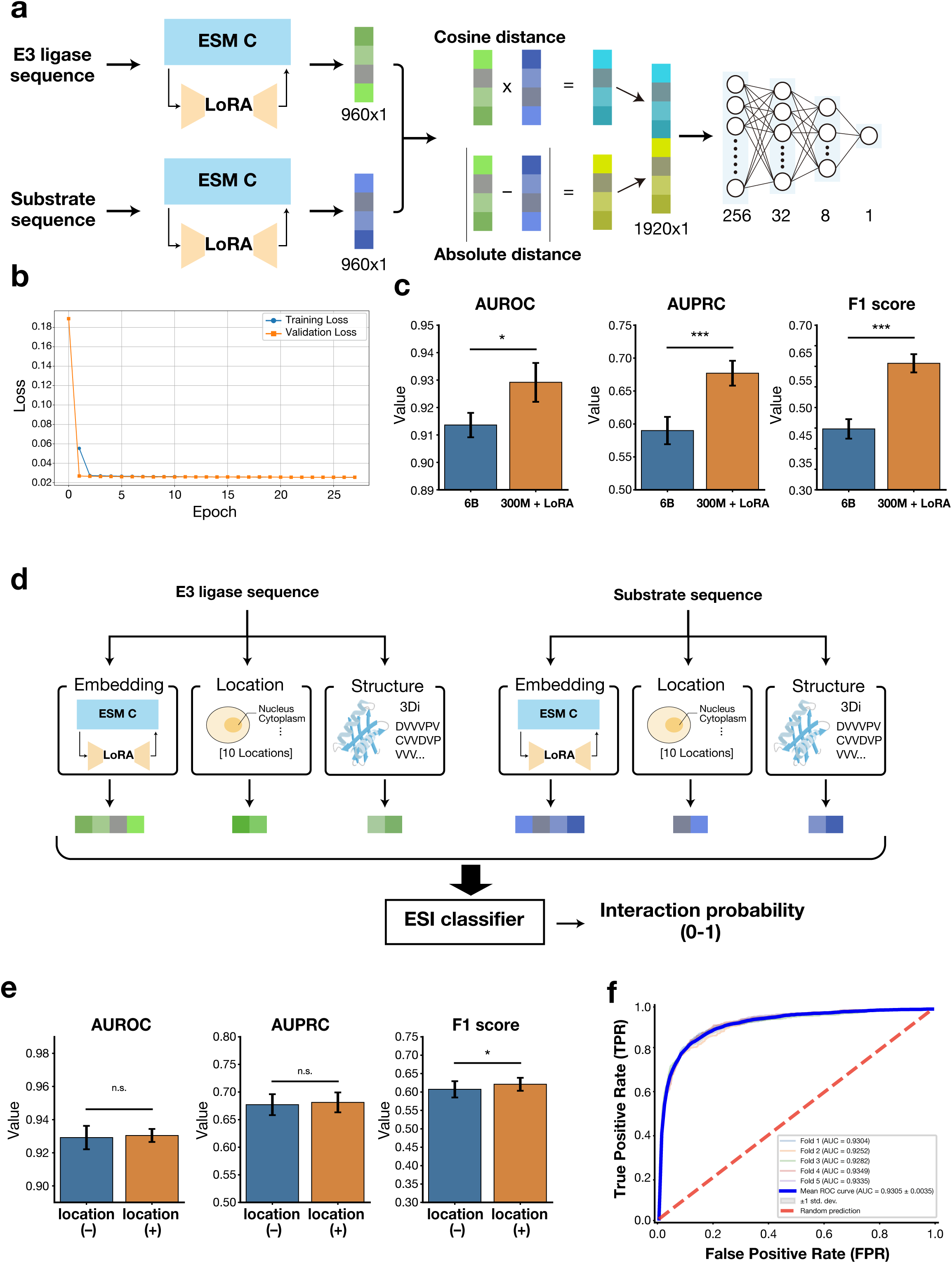
Task-adapted protein language models and multimodal integration drive state-of-the-art ESI prediction by Ubicon. (**a**) Schematic of the contrastive learning architecture employing PEFT via LoRA. This approach adapts pre-trained ESM-C 300M embeddings specifically for the ESI recognition task, using distances between paired E3-substrate representations as learning signals. (**b**) Representative learning curves demonstrate stable convergence during LoRA fine-tuning without evidence of overfitting, validating the training procedure. Training loss (blue) and validation loss (orange) are shown. (**c**) Task adaptation dramatically boosts predictive power. Comparison reveals that the lightweight PLM adapted via LoRA (300M + LoRA, combined with 3Di features) significantly and substantially outperforms the much larger, general pre-trained PLM (6B, combined with 3Di features) across all key metrics: AUROC, AUPRC, and F1 score (paired *t*-test; \*\*\**p* < 0.001, \**p* < 0.05). This striking result underscores the power of targeted, efficient model specialization over sheer scale for this complex biological task. Error bars indicate SD (n=5 folds). (**d**) Architecture of the final Ubicon predictor. Ubicon leverages a multimodal approach, integrating task-adapted PLM embeddings, predicted structural features (3Di), and predicted subcellular localization information as input to ESI classifier, which outputs the ESI probability (0-1). Created with BioRender.com. (**e**) Incorporating cellular context enhances prediction accuracy. Adding predicted subcellular localization features (location (+)) to the LoRA-adapted PLM and structure features (location (-)) yields a statistically significant improvement in the F1 score (paired *t*-test; \**p* < 0.05), highlighting the importance of cellular context for balancing precision and recall, while AUROC and AUPRC remain high and statistically unchanged (n.s.). Error bars indicate SD (n=5 folds). (**f**) Ubicon achieves outstanding discriminative performance. Receiver operating characteristic (ROC) curves from rigorous 5-fold cross-validation of the final Ubicon model. The mean Area Under the Curve (AUC) reaches 0.9305 ± 0.0035 (mean ± SD), demonstrating state-of-the-art performance significantly exceeding random prediction (dashed red line). Individual fold curves are shown alongside the mean curve (bold blue).

Next, building on the principle that interacting proteins must co-localize within the cell^44,45^, we incorporated subcellular localization predictions as a contextual filter. We added predicted localization probabilities across 10 compartments (via DeepLoc2^46^) for both E3 and substrate as input features (Fig. 2d). This approach is biologically sound, as confirmed by high predicted co-localization probabilities for known interacting pairs like nuclear MDM2-p53^47^ and nuclear/cytoplasmic Fbxw7-c-Myc^48^ (Fig. S2a, b). Integrating localization features significantly improved the LoRA-tuned model’s F1 score, primarily by increasing precision through the effective filtering of biologically implausible interactions (Fig. 2e). This demonstrates the synergistic benefit of combining molecular features (sequence, structure) with cellular context (localization) for accurate *in vivo* interaction prediction.

The sequential integration of task-adapted embeddings, structural features, and localization context culminated in our final prediction framework, Ubicon (Table S1). Rigorous 5-fold cross-validation demonstrated Ubicon’s superior predictive accuracy, achieving a mean Area Under the Receiver Operating Characteristic curve (AUROC) of 0.9305 and a mean Area Under the Precision-Recall curve (AUPRC) of 0.6812. The high AUPRC is particularly noteworthy, indicating Ubicon’s strong ability to precisely identify true positive interactions within the imbalanced ESI landscape, a critical feature for biological discovery. Crucially, Ubicon significantly outperforms existing state-of-the-art ESI prediction methods. For instance, its AUPRC of 0.6812 represents a substantial improvement over the previous leading method, DeepUSI (AUPRC 0.4445) (Table S1), highlighting its enhanced capability to reduce false positives and reliably identify genuine enzyme-substrate interactions.

To enhance Ubicon’s utility for downstream applications, such as prioritizing candidates for experimental validation, we calibrated its output scores to better reflect true interaction probabilities. Raw prediction scores from complex models can exhibit biases; therefore, we evaluated isotonic regression^49^ and temperature scaling^50^ for calibration, finding isotonic regression yielded the best results (Fig. S3a). Based on this comparative analysis, we adopted isotonic regression as our calibration method of choice. Applying this method effectively aligned Ubicon’s predicted scores with empirical interaction frequencies across the score range. We refer to these calibrated predictions as "Ubicon scores", which represent the model’s estimate of E3-substrate interaction probability on a scale from 0 to 1. This calibration step provides trustworthy, interpretable probabilities—for example, a calibrated Ubicon score of 0.8 reliably signifies an ∼80% likelihood of a genuine interaction—enabling researchers to set meaningful confidence thresholds for subsequent studies (Fig. S3b, c).

### Ubicon accurately recapitulates known biology and deciphers functional determinants of ESI recognition

Beyond predictive accuracy, a critical challenge for computational models like Ubicon is whether they exhibit genuine biological understanding. We therefore interrogated Ubicon’s ability to recapitulate established biological knowledge and capture molecular determinants governing E3-substrate interactions through multifaceted *in silico* validation.

First, we confirmed Ubicon’s capacity to recognize diverse, experimentally validated ESIs. The model consistently assigned high-confidence scores much greater than 0.5 to benchmark pairs fundamental to cell regulation, such as FBXL5-IRP2^51,52^ and VHL-HIF1α^53^ (Fig. 3a).

**Figure 3.**
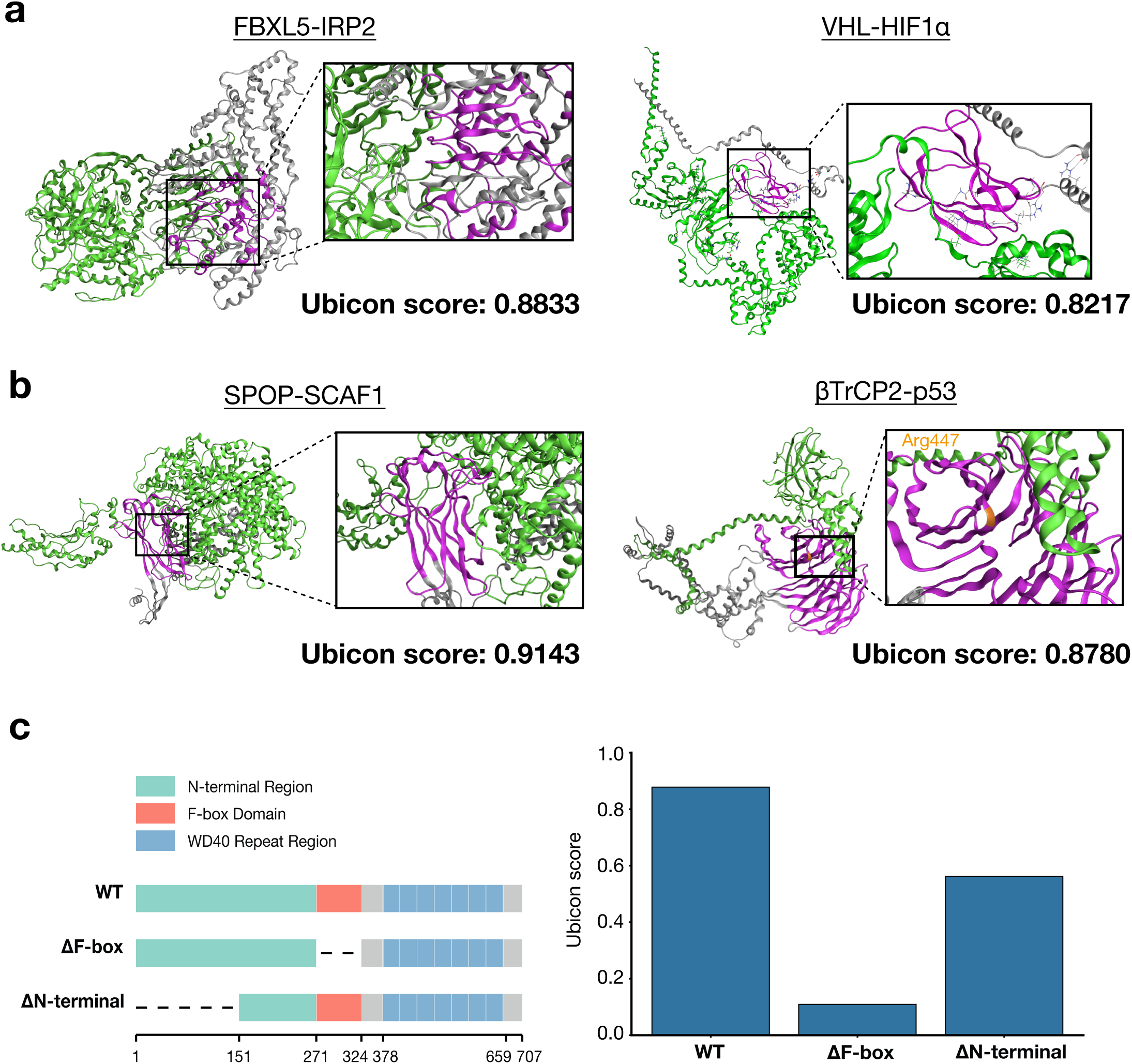
Ubicon captures the molecular basis and functional determinants of E3-substrate recognition. (**a**) Accurate recapitulation of canonical ESI interactions. Ubicon correctly predicts high interaction probabilities (Ubicon scores) for well-established E3-substrate pairs crucial for cellular homeostasis and disease, exemplified by FBXL5-IRP2 (left) and VHL-HIF1α (right). Structural modeling corroborated these predictions, indicating consistency with known binding modes and engagement of critical interaction interfaces. (**b**) Robust generalization demonstrates learning of interaction principles. Ubicon exhibits strong generalization capabilities, accurately predicting interactions absent from its training dataset with high confidence, as shown for independent test cases SPOP-SCAF1 (left) and βTrCP2-p53 (right). Structural modeling further supports the biological plausibility of these predictions, suggesting interactions mediated via established recognition domains (e.g., the MATH domain of SPOP; the WD40 repeats of βTrCP2) and potentially involving residues experimentally verified as critical for binding (e.g., near Arg447 in βTrCP2). (**c**) Dissecting domain function via *in silico* mutagenesis reveals Ubicon’s mechanistic understanding. To rigorously test if Ubicon comprehends the functional significance of specific protein domains in mediating ESIs, we performed *in silico* deletion analyses using the FBXW7-c-Myc interaction as a paradigm. Left: Domain architecture of FBXW7 showing regions targeted for deletion (ΔF-box, ΔN-terminal). Right: Ubicon-predicted interaction probabilities. Deletion of the F-box domain (ΔF-box), known to be indispensable for substrate recognition, completely abrogates the predicted interaction, precisely mirroring its biological essentiality. Strikingly, deletion of the larger N-terminal domain (ΔN-terminal), which is dispensable for c-Myc binding due to functional isoforms, results in a maintained high interaction score. This stark contrast provides compelling evidence that Ubicon learns the hierarchical functional importance of distinct domains with remarkable accuracy. This capability signifies that Ubicon captures biologically meaningful features far beyond statistical patterns, establishing its potential to provide mechanistic insights into ESI determination.

Next, we assessed Ubicon’s generalization capability on interactions absent from the original data as external validation study. Ubicon predicted recently identified ESIs, including SPOP-SCAF1^25^ and β-TrCP2-p53^54^, with high confidence scores (Fig. 3b). Structural models for these predictions again suggested biologically plausible binding modes involving canonical recognition domains. This robust performance on unseen data indicates that Ubicon captures generalizable molecular principles governing ESI specificity, supporting its potential for novel discovery.

To probe if Ubicon grasps the functional importance of specific protein domains critical for mediating ubiquitination, we performed *in silico* domain deletion experiments using the well-characterized Fbxw7-c-Myc interaction. Deleting the F-box domain of Fbxw7—which is essential for recruiting the core SCF complex components (Skp1-Cul1-Rbx1) needed for ubiquitination — reduced the Ubicon score to near zero (Fig. 3c). This mirrors the biological outcome where F-box deletion prevents c-Myc ubiquitination even if substrate binding persists^55^, strongly suggesting the Ubicon score reflects the capacity for functional ubiquitination rather than merely physical binding. In striking contrast, deleting the large N-terminal region of Fbxw7— known to be dispensable for core SCF complex formation^56^—resulted in only a modest reduction in the Ubicon score. This differential sensitivity provides compelling evidence that Ubicon learns functional context, distinguishing critical domains from less essential regions for specific interactions.

The model’s ability to correctly predict the differential impact of these deletions suggests it captures nuanced structure-function relationships. This capacity indicates Ubicon operates beyond pattern recognition, offering potential for mechanistic insights into E3-substrate recognition and serving as a reliable pipeline for investigating ESI biology.

### Ubicon captures deeply conserved principles of E3-Substrate recognition across eukaryotic evolution

A pivotal question is whether the molecular principles governing E3-substrate recognition are conserved across vast evolutionary distances^57,58^. We investigated if Ubicon—trained primarily on human data but incorporating evolutionary information via PLMs and structural constraints—could generalize to decipher such rules. First, evaluating Ubicon in *Mus musculus*, we found it accurately identified known, functionally conserved orthologous ESIs (e.g., Mdm2-p53^59^, Fbxw7-Myc^56^, RNF4-PARP1^60^) with high confidence scores comparable to human predictions (Table S2). Furthermore, predicted structural models for mouse ESI complexes revealed plausible interfaces consistent with conserved recognition mechanisms (Fig. S4a). This strong performance in a closely related species provided initial evidence that Ubicon captures evolutionarily conserved recognition principles.

Building on this, we assessed Ubicon’s generalization capacity across deeper evolutionary divides using curated ESI datasets from diverse eukaryotic kingdoms: animals (*D. melanogaster*, *C. elegans*), fungi (*S. cerevisiae*), and plants (*A. thaliana*). Remarkably, Ubicon successfully assigned high scores (Ubicon score > 0.5) to a substantial fraction of known ESIs within each lineage, demonstrating significant predictive capability despite the vast phylogenetic distances (Fig. S4b). This ability to recognize conserved interaction features across kingdoms was further exemplified by the accurate, high-confidence prediction of specific orthologous interactions, such as the *Drosophila* rdx-CG2926 pair (homologous to human SPOP-CCNB1), supported by structural modeling (Fig. S4c). Taken together, Ubicon’s robust performance across this broad phylogenetic range indicates it has learned deeply conserved molecular features governing ESI specificity throughout eukaryotic evolution.

### Systems analysis of the high-resolution human “ESIome” unveils architectural principles and functional organization

Leveraging the optimized and calibrated Ubicon framework, we performed systematic E3-substrate interaction predictions across the human proteome, evaluating potential interactions between E3 ligases and approximately 20,000 cellular proteins. This extensive computation generated high-confidence interaction probabilities for nearly six million potential ESI pairs, culminating in the comprehensive, high-resolution predictive atlas of the human ESI network— termed the "ESIome" (Table S3). This unprecedented resource provided the foundation to interrogate the global architecture, design principles, and functional logic of this critical regulatory network at a systems level.

We initiated our exploration by constructing the ESI network graph using high-confidence predictions (Ubicon score ≥ 0.8), resulting in a complex interactome comprising 207,816 predicted interactions connecting 486 E3 ligases with nearly 20,000 substrate proteins (Fig. 4a). Initial topological analysis revealed network properties markedly distinct from those of random networks. The calculated average node degree (25.85) and high average clustering coefficient (0.5861) strongly indicated a non-random, intrinsically organized structure^61^. This finding suggests the human ESIome is a highly structured system shaped by specific biological constraints, prompting deeper investigation into its organizational principles.

**Figure 4.**
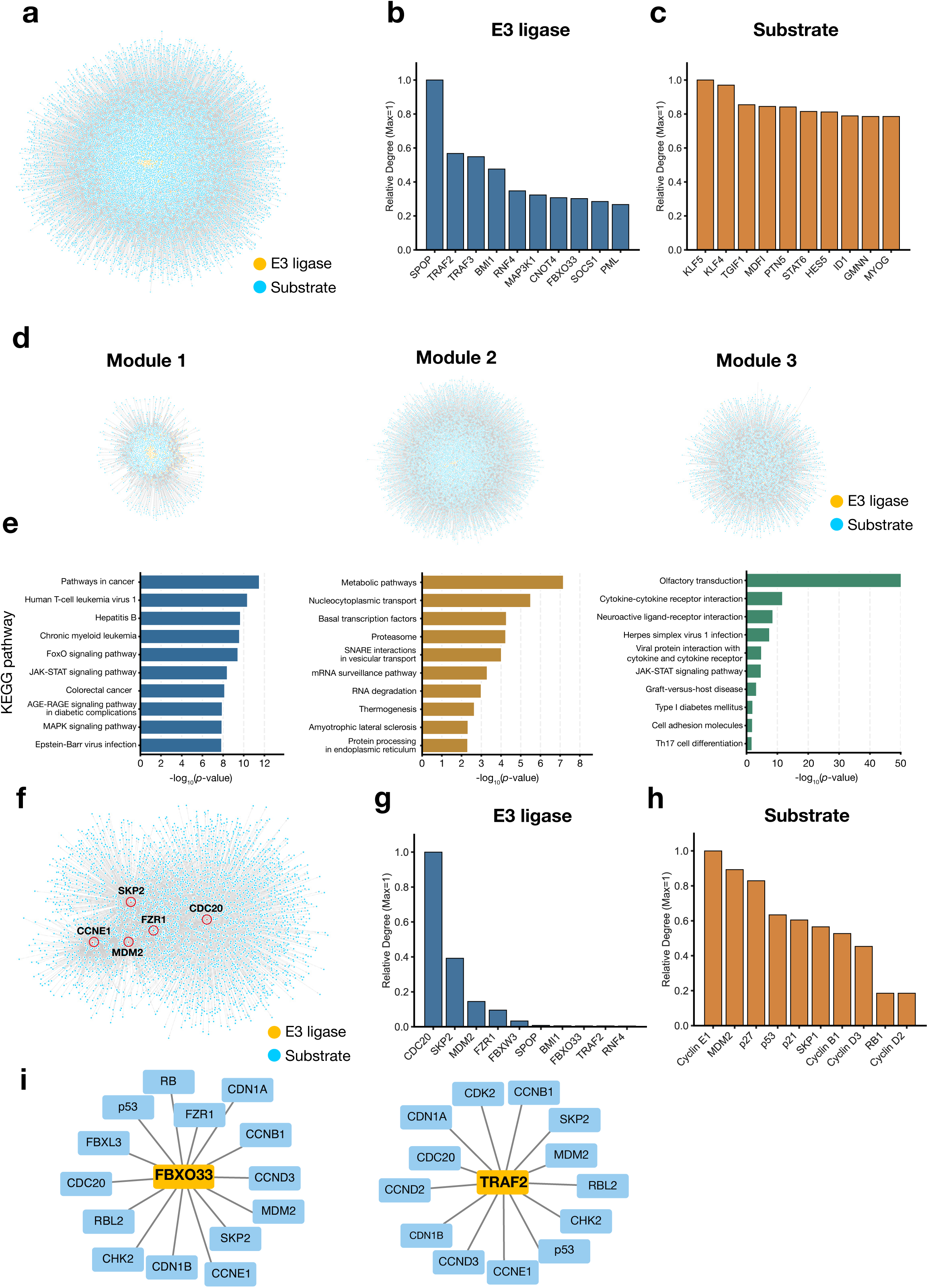
Mapping the human E3 ligase-substrate interactome defines its functional modules and regulatory hubs. (**a)** Global view of the predicted ESI network (“ESIome”) across the human proteome. Nodes represent E3 ligases (orange) and substrate proteins (blue), while edges indicate predicted interactions. The ESIome was visualized using Cytoscape, and the layout was optimized using the Prefuse Force Directed algorithm. (**b)** Top 10 hub E3 ligases ranked by degree (number of interacting substrates). The bar graph shows the relative number of substrates interacting with each E3 ligase, identifying potential key regulators controlling numerous substrates across the proteome. (**c)** Top 10 hub substrate proteins ranked by degree (number of interacting E3 ligases). The bar graph shows the relative number of E3 ligases interacting with each substrate, suggesting these are key targets regulated by multiple E3s. (**d)** Network community (module) detection using the Louvain method. Visualization by Cytoscape reveals three major modules composed of densely interconnected protein groups, suggesting the clustering of proteins involved in specific biological functions. (**e)** KEGG pathway enrichment analysis for each community identified in (d). The bar graph displays significantly enriched pathways within each community. The y-axis shows pathway names, and the x-axis represents the statistical significance of enrichment (-log_10_(*p*-value)). Pathways related to cancer (Module 1), metabolism (Module 2), and olfactory transduction/intercellular signaling (Module 3) were significantly enriched. (**f)** Subnetwork constructed by extracting proteins (E3 ligases and substrates) associated with a specific biological process (cell cycle). Nodes are colored as in (a) (E3 ligases: orange; substrates: blue), and the layout was generated using the Prefuse Force Directed algorithm. Key hub proteins involved in cell cycle regulation are highlighted. (**g)** Relative degree of key E3 ligases within the cell cycle subnetwork. The bar chart quantifies the connectivity of central E3s, including canonical regulators (e.g., CDC20, SKP2, MDM2) and other prominent hubs identified in this context (e.g., FBXO33, TRAF2). (**h)** Relative degree of key substrates within the cell cycle subnetwork. The bar chart quantifies connectivity, highlighting prominent substrate hubs such as Cyclin E1 (CCNE1). The high degree of MDM2 as a substrate underscores its dual E3/substrate role, indicative of feedback regulation. (**i)** Subnetwork interactions centered on two newly identified hubs in cell cycle regulation. Left: FBXO33 interacts with multiple critical cell cycle regulators including p53, RB1, CDC20, and several cyclins. Right: TRAF2 similarly shows extensive connections with key cell cycle proteins, suggesting these E3 ligases may play previously unrecognized roles in cell cycle control.

Recognizing that the structure and function of many complex biological networks are often governed by highly connected hub nodes ^62,63^, we performed centrality analysis to identify key regulatory entities within the ESIome. Node degree analysis pinpointed several E3 ligases as prominent hubs, including known master regulators frequently implicated in cancer such as SPOP^64,65^, TRAF2^66^, and TRAF3^67,68^, each predicted to interact with a disproportionately large number of substrates (Fig. 4b). Consistent with their established biological roles, these hub E3s participate broadly in processes like cell proliferation, apoptosis, and DNA repair. Concurrently, key transcription factors crucial for cell fate determination (e.g., KLF4^69^, KLF5^70^, TGIF1^71^) and critical regulators like STAT6^72^ (immune response), and GMNN^73^ (DNA replication) emerged as major substrate hubs targeted by numerous E3s (Fig. 4c). These observations strongly suggest that the ESI network architecture exhibits scale-free characteristics, organized around a limited set of functionally critical E3 and substrate hubs.

A fundamental question in systems biology concerns the functional organization of complex networks into specialized modules^62,63^. We probed this principle within the predicted ESI network using community detection algorithms coupled with pathway enrichment analysis. Application of the Louvain algorithm^74^ robustly partitioned the ESI network into three major, densely interconnected modules (Fig. 4d). Strikingly, KEGG pathway enrichment analysis of the substrates within each module revealed a remarkable degree of functional specialization (Fig. 4e). Module 1 substrates were overwhelmingly enriched in pathways related to cancer signaling, viral responses, and cell fate control. Module 2 predominantly comprised proteins involved in core cellular machinery, including metabolism, transport, and protein homeostasis pathways. Module 3 displayed extreme enrichment for sensory perception (particularly olfaction), immune system processes, and intercellular communication pathways. This analysis uncovers a pronounced functional segregation within the human ESIome, revealing its organization into highly specialized modules governing distinct cellular realms—cancer/cell fate, core metabolism/housekeeping, and sensory/immune functions—likely facilitating efficient domain-specific regulation and controlled crosstalk.

We then assessed Ubicon’s ability to dissect specific biological circuits using cell cycle control as a test case. The extracted cell cycle subnetwork (Fig. 4f) accurately recapitulated known biology, identifying canonical E3 hubs (e.g., APC/C-CDC20^75,76^, SCF-SKP2^77^, MDM2^78^ and substrate hubs (e.g., Cyclin E1) (Fig. 4g, h), including regulatory motifs like MDM2’s dual E3/substrate role^79,80^ (Fig. 4h). Importantly, the analysis extended beyond established knowledge, identifying E3s like FBXO33 and TRAF2^66^ as unexpectedly central nodes in this context (Fig. 4i). Notably, Ubicon predicted the FBXO33-p53 interaction before its recent experimental confirmation^81^, despite this interaction being absent from our training data, thus validating Ubicon’s predictive power. This demonstrates that the Ubicon-derived ESI network serves as a powerful discovery engine, capable of identifying novel regulatory links prior to experimental validation.

Collectively, these systems-level analyses, enabled by the predictive power of Ubicon, provide an unprecedented multi-scale perspective on the human ESIome—from its global topology and central hubs to its functional modularity and detailed circuit wiring. The elucidated architectural principles and novel predictions furnish numerous avenues for future experimental investigation, significantly advancing our understanding of ubiquitin-mediated regulation.

### Ubicon deciphers cancer-specific ESI network rewiring linked to oncogenic pathways and clinical outcomes

Dysregulation of the ubiquitin system is a hallmark of cancer^4,5^, yet how specific ESI network perturbations contribute to tumorigenesis remains poorly understood. To bridge this critical knowledge gap, we integrated our high-fidelity ESI classifier, Ubicon, with multi-omics data from The Cancer Genome Atlas^82^, seeking to delineate the functional consequences of altered E3 ligase expression in human cancers (Fig. 5a). We first aimed to pinpoint E3 ligases whose aberrant expression profiles might represent key nodes driving ESI network remodeling across diverse malignancies. Pan-cancer transcriptomic analysis revealed distinct E3 ligase clusters exhibiting consistent upregulation or downregulation across multiple tumor types (Fig. 5b). Focusing on breast invasive carcinoma (BRCA) and lung adenocarcinoma (LUAD) as representative examples, we identified a cohort of E3s significantly overexpressed in tumors relative to adjacent normal tissues. From the E3s demonstrating the most statistically significant upregulation (log_2_ fold change > 1, adjusted *p*-value < 0.01), we selected UHRF1, AURKA, and CDC20 for further investigation, specifically prioritizing them as they were among the most significant E3 ligases. These E3s are master regulators of genome stability and cell cycle progression whose functions are frequently commandeered during malignant transformation^83–86^. These findings implicate specific, functionally critical E3 ligases as putative oncogenic drivers whose transcriptional upregulation may initiate widespread perturbations within the ESI network in these cancers.

**Figure 5.**
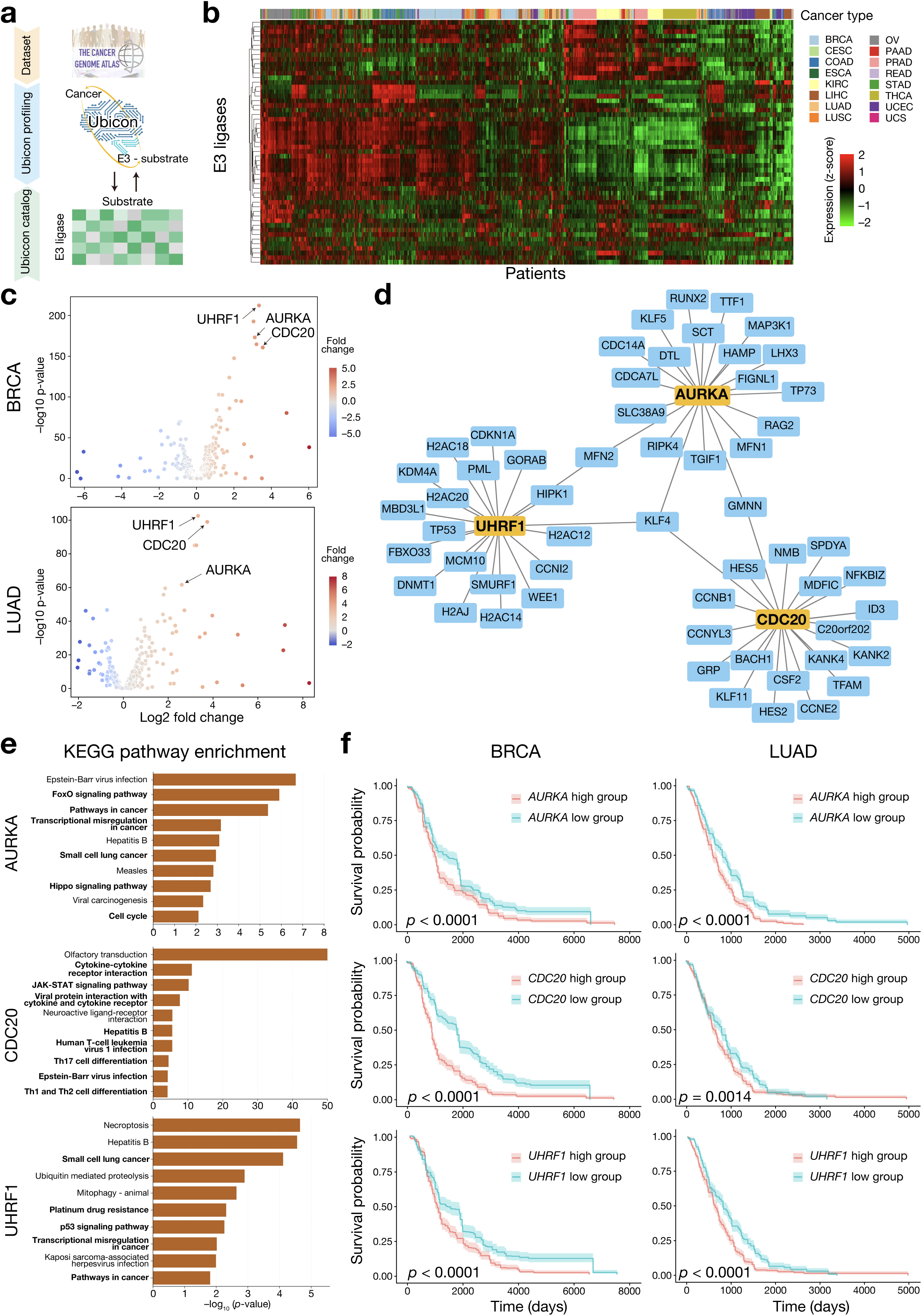
Systems analysis reveals E3 ligase network aberrations across human cancers. (**a**) Workflow integrating Ubicon ESI predictions with TCGA datasets for pan-cancer analysis of E3-substrate network alterations. Created with BioRender.com. (**b**) Pan-cancer E3 ligase expression profiles in TCGA tumors. Heatmap shows mRNA expression z-scores, clustered hierarchically by E3s (rows) and samples (columns; cancer types indicated in legend). Red indicates high expression, green low expression. (**c**) Differential expression of E3 ligases in breast cancer (BRCA, left) and lung adenocarcinoma (LUAD, right). Volcano plots show log_2_ fold change (tumor vs. normal) versus statistical significance (-log_10_ adjusted *p*-value; Wald test, Bonferroni). Red/blue dots indicate significant upregulation/downregulation. Commonly dysregulated E3s (AURKA, CDC20, UHRF1) are labeled. (**d**) Predicted interaction network for key dysregulated E3s identified in (c) (AURKA, CDC20, UHRF1; orange) and their high-confidence substrate candidates. Network visualized using Cytoscape. (**e**) KEGG pathway enrichment analysis of predicted substrates (d) for AURKA, CDC20, and UHRF1. Bar plots show significantly enriched pathways (-log_10_ *p*-value). (**f**) Kaplan-Meier analysis correlating *AURKA*, *CDC20*, and *UHRF1* mRNA expression with patient survival in BRCA and LUAD (TCGA). Patients stratified by median expression into high (red line) and low (blue line) expression groups. *P*-values determined by log-rank test.

To elucidate the mechanistic links between E3 dysregulation and cancer phenotypes, we employed Ubicon to predict the substrate networks governed by the prominently upregulated E3s: UHRF1, AURKA, and CDC20. Ubicon delineated distinct sets of high-confidence substrates for each E3 (Fig. 5d), providing a putative map of the downstream signaling pathways affected by their overexpression. Subsequent pathway enrichment analysis of these predicted substrate networks revealed striking functional specificities (Fig. 5e). The predicted UHRF1 substrate network was significantly enriched for proteins involved in DNA replication and chromatin regulation, aligning precisely with its established role in maintaining genome integrity^87,88^. In contrast, the predicted AURKA substrate network converged on pathways governing cell proliferation, survival, and oncogenic signaling cascades. Distinctly again, the predicted CDC20 substrates were strongly enriched in pathways related to immune modulation, including T cell differentiation and cytokine receptor signaling. These computationally derived insights suggest a model where the upregulation of specific E3s contributes to core cancer hallmarks by perturbing distinct functional modules: UHRF1 potentially driving genomic instability, AURKA promoting uncontrolled proliferation and survival, and CDC20 possibly modulating the tumor immune microenvironment.

Finally, to ascertain the clinical relevance of these specific E3 upregulations and the associated ESI network alterations predicted by Ubicon, we rigorously assessed their correlation with patient survival outcomes. Strikingly, elevated transcript levels of UHRF1, AURKA, or CDC20 each demonstrated a significant and robust association with poorer overall survival in both BRCA and LUAD patient cohorts from TCGA (Fig. 5f). This robust clinical correlation provides compelling evidence for the prognostic significance and likely functional importance of these E3 ligases in cancer progression. Furthermore, it strongly suggests that the ESI network rewiring downstream of these E3s, as predicted by Ubicon, directly contributes to tumor aggressiveness and shapes patient prognosis. Collectively, our integrated analysis, powered by Ubicon, reveals a systems-level mechanism connecting E3 ligase upregulation to specific ESI network perturbations, consequent oncogenic pathway dysregulation, and adverse clinical outcomes, thereby highlighting these E3s and their downstream networks as potential therapeutic targets.

## Discussion

In this study, we developed Ubicon, a deep learning framework that integrates multimodal biological data to predict E3 ubiquitin ligase-substrate interactions (ESIs) with high accuracy. Ubicon addresses the long-standing challenge of comprehensively mapping the human ESI network by effectively combining information from task-adapted protein language models (PLMs), predicted structures, and subcellular localization context. Our work establishes a new state-of-the-art in ESI prediction (AUROC=0.9305, AUPRC=0.6812), provides critical insights into the biological determinants of these interactions through *in silico* validation, generates the first high-resolution predictive atlas of the human ESIome, and uncovers cancer-specific network alterations linked to clinical outcomes.

Ubicon’s performance surpasses previous computational methods, which often relied on limited features like sequence motifs or domain data, thereby struggling with novel interactions or complex recognition modes. By integrating richer sequence context, structural information, and cellular localization, Ubicon achieves high-confidence identification of true ESIs within the vast search space of the proteome. This capability can significantly accelerate ubiquitin research by enabling more focused, hypothesis-driven experimental validation strategies, moving beyond the limitations of traditional large-scale screens.

The technical advance underpinning Ubicon lies in the synergistic integration of diverse data types, critically optimized by task-specific adaptation of PLMs. We demonstrated that PEFT via LoRA^35^ allowed a moderately sized PLM (ESM-C 300M) to outperform a much larger generalist model (ESM-C 6B) specifically for ESI recognition (Fig. 2c). This highlights the power of efficiently adapting large pre-trained models for specific biological tasks and establishes the combination of adapted sequence embeddings, 3Di structural states, and localization data as a potent feature set for predicting molecular interactions (Table S1).

Our systems-level analysis of the Ubicon-predicted ESIome offers key insights into the organizational principles of ubiquitin-mediated regulation. We identified central E3 (e.g., SPOP, TRAF2/3) and substrate (e.g., KLF4/5, GMNN) hubs that likely serve as critical control points in cellular networks, potentially representing vulnerabilities in disease states (Fig. 4). Furthermore, the ESI network exhibits clear functional modularity, with distinct modules enriched for specific biological processes like cancer signaling, core metabolism, or sensory/immune functions (Fig. 4e). This modular architecture provides a framework for understanding how specific regulatory logic is implemented within the broader "ubiquitin code".

Integrating Ubicon predictions with TCGA cancer genomics data provided direct evidence for the pathological relevance of ESI network perturbations. Specific E3s frequently upregulated in breast and lung cancer (AURKA, CDC20, UHRF1) were linked, via their Ubicon-predicted substrate networks, to the dysregulation of core oncogenic pathways (cell cycle, genome maintenance) and significantly correlated with poor patient survival (Fig. 5). This analysis not only deepens our mechanistic understanding of how E3 dysregulation drives cancer but also yields critical translational insights. The ability of Ubicon to map E3-substrate specificity with high resolution has potential to advance targeted protein degradation (TPD) therapies^13,14^. For instance, Ubicon can inform the selection of E3 ligases to recruit for degrading specific targets (e.g., via PROTACs), predict the substrate profiles (including potential off-targets) of chosen E3s, and help evaluate how mutations might impact degrader efficacy (Fig. 3c), thereby accelerating the rational design and optimization of TPD strategies.

While Ubicon represents a significant advance, limitations remain. Predictions can be influenced by biases in current training data, and performance may vary for understudied E3s. Future iterations will benefit from incorporating newly validated interactions. The model relies primarily on static predicted structures and does not explicitly account for protein dynamics, complex formation effects, or the influence of other post-translational modifications, representing key areas for future development. Furthermore, integrating dynamic cellular context beyond localization, such as protein abundance or cell state information from multi-omics data, could further refine prediction accuracy.

In conclusion, Ubicon establishes a new benchmark for ESI prediction by effectively integrating task-adapted PLMs, structure, and localization data. It moves beyond prediction accuracy to provide biological insights, reveal systems-level network organization, and connect ESI perturbations to disease states and clinical outcomes. The comprehensive ESIome atlas generated by Ubicon (Table S3) serves as a valuable community resource. We anticipate that Ubicon and its underlying multimodal integration strategy will catalyze discoveries in ubiquitin biology and accelerate the development of novel therapeutic strategies targeting various diseases.

## Materials and Methods

### Dataset construction

All protein sequences used in this study were sourced from UniProt Knowledgebase (UniProtKB)^89^. Sequences exceeding 2046 amino acids were excluded due to the input length constraints of the ESM-C model employed for generating sequence embeddings^36^.

For training and validation, the positive dataset, comprising known human E3-substrate interaction (ESI) pairs, was compiled based on curated enzyme-substrate relationships documented in the UbiBrowser 2.0 database^27^ (downloaded on February 6, 2025). This initial set was refined through manual curation to exclude pairs where the designated enzyme was not a confirmed or putative E3 ligase according to established E3 catalogues or literature evidence. This stringent filtering process yielded a high-confidence positive dataset of 2,727 unique ESI pairs. The negative dataset was designed to represent protein pairs that physically interact but lack known E3-substrate relationships. To construct this set, human protein-protein interactions (PPIs) involving at least one known or putative E3 ligase were extracted from the BioGrid database^90^ (version 4.4, accessed on February 15, 2025). Any pairs identical to those included in the positive dataset were subsequently removed from the extracted PPIs. This procedure resulted in a negative dataset of 28,125 protein pairs.

### Feature Extraction

#### Sequence Embeddings

Per–residue sequence embeddings were generated using pre-trained protein language models (PLMs). Specifically, embeddings from ESM-C (6B parameters) were obtained via the official Evolutionary Scale API^36^. For local computation and fine-tuning, embeddings were generated using the ESM2 model^91^ (15B parameters, HuggingFace model facebook/esm2_t48_15B_UR50D) and the ESM-C model (300M parameters, HuggingFace model Synthyra/ESMplusplus_small^92^, approximating ESM Cambrian 300M). For input to downstream models requiring fixed-size protein representations, per-protein embeddings were derived by averaging the corresponding per-residue embeddings.

### LoRA Fine-tuning of ESM-C 300M

To adapt PLM representations specifically for ESI prediction, we applied parameter-efficient fine-tuning (PEFT) using Low-Rank Adaptation (LoRA)^35^ to the ESM-C 300M model. LoRA modules (rank r=4, dropout=0.1, lora_alpha=32) were inserted into the final 20 attention blocks of the PLM. Fine-tuning was performed using a contrastive learning approach. The downstream classification head used during LoRA training consisted of a 3-layer multilayer perceptron (MLP) with architecture [256, 32, 8]. This MLP took as input a vector derived from concatenating the cosine similarity and absolute difference between the E3 ligase and substrate embeddings generated by the LoRA-adapted ESM-C model (Fig. 2a). The same set of LoRA parameters was applied to generate both the E3 and substrate embeddings within this fine-tuning procedure. Model construction and training were implemented using PyTorch (version 2.1.0).

### Structural Features

Three-dimensional structural information was incorporated by representing protein structures as 1D discretized structural state sequences (3Di sequences) using Foldseek^37^. Structural data were primarily obtained from the AlphaFold Protein Structure Database (AlphaFold DB)^93^. For proteins lacking an entry in AlphaFold DB, structures were predicted *de novo* using AlphaFold2^34^. The foldseek *createdb* command was used to generate 3Di state sequences from PDB-formatted structure files for use as model input features.

### Subcellular Localization Features

Predicted subcellular localization probabilities were used as contextual features. For each protein sequence, the probability of localization to 10 distinct cellular compartments (Cytoplasm, Nucleus, Extracellular, Cell membrane, Mitochondrion, Plastid, Endoplasmic reticulum, Lysosome/Vacuole, Golgi apparatus, Peroxisome) was predicted using DeepLoc2^46^ in ‘fast mode’. These 10 probability values were concatenated and used directly as input features to the prediction model.

### Interaction Prediction Models

#### Baseline CNN Model

A baseline Convolutional Neural Network (CNN) model was constructed, inspired by architectures like DeepUSI and those available in DeepPurpose^94^, to predict E3-substrate interactions (ESIs). This model accepted E3 ligase and substrate inputs represented by their respective amino acid sequences. These sequences were processed to extract features: fixed-length protein embeddings obtained by averaging per-residue embeddings from ESM-C (6B, without LoRA), and structural features derived from 3Di sequences (embedded into 64 dimensions using torch.nn.Embedding). Each protein’s sequence embedding and structural feature vector were processed through separate blocks of 1D CNN layers with varying kernel sizes (e.g., 3, 4, 5) designed to capture local patterns. Max-pooling layers were applied after convolutional blocks to obtain fixed-length representations. The resulting latent vectors for the E3 and substrate were concatenated and fed into a Multi-Layer Perceptron (MLP) classifier consisting of fully connected layers with ReLU activations and dropout. The final layer used a sigmoid activation function to output an interaction probability between 0 and 1. The model was trained using a Binary Cross-Entropy loss function with the Adam optimizer. This CNN served as a baseline for comparison against subsequent models.

### Tree-based and Tabular Models

To identify the optimal architecture for classifying ESIs based on our final feature set, we implemented and evaluated several decision tree-based models and other established methods for tabular data. The decision tree based models included, CatBoost^38^, XGBoost^39^, and LightGBM^40^alongside Random Forest^41^. We also evaluated deep learning models specialized for tabular data: TabNet^42^ and the pre-trained transformer model TabPFN^43^. Input features for these models consisted of a concatenated vector comprising: (i) the mean protein embedding derived from the LoRA-tuned ESM-C 300M model, (ii) 3Di frequency information: a vector representing the distribution of a subset of structural encoding characters in the protein’s 3D representation. This was derived by calculating the relative frequencies of specific characters in the 3Di sequence and normalizing to capture the structural signature independent of protein length, and (iii) the 10-dimensional subcellular localization probability vector.

Hyperparameters for each model (excluding the TabNet and TabPFN) were systematically optimized using the Optuna framework^95^ to maximize the mean AUROC during 5-fold cross-validation on the training dataset. Key hyperparameters tuned typically included parameters controlling tree complexity (e.g., depth, number of leaves), learning rates, regularization terms, and feature/data subsampling rates (Table S4). Model implementations utilized the official libraries for CatBoost (version 1.2.7), LightGBM (version 4.6.0), XGBoost (version 2.1.4), scikit-learn (version 1.3.2), the pytorch-tabnet library (version 4.1.0) for TabNet, and the tabpfn library (version 0.1.11) for TabPFN.

### Comparison with Other E3-Substrate Prediction Methods

To benchmark against existing approaches, we evaluated two alternative methods for E3-substrate interaction prediction. First, we tested the DeepPurpose library as previously reported^94^, representing a standard implementation similar to previously published methods like DeepUSI^28^. We employed the default CNN-based interaction prediction model within DeepPurpose, providing only the E3 ligase and substrate amino acid sequences as input. Hyperparameters for the DeepPurpose CNN model, including MLP architecture (layer count, neuron count), CNN architecture (layer count, filter numbers, kernel sizes), learning rate, batch size, and dropout rates, were optimized using Optuna^95^ to maximize AUROC on our dataset through 5-fold cross-validation (Table S4).

Additionally, we evaluated E3-targetPred as a second alternative deep learning approach for predicting E3-substrate interactions^29^. E3-targetPred utilizes a neural network approach based on CKSAAP feature encoding. Since our attempts to retrain the E3-targetPred model on our dataset resulted in model convergence issues where all test samples were predicted as negative, we used the model trained on the authors’ original training dataset to evaluate its performance on our curated E3-substrate pairs. These approaches allowed us to assess the generalizability of both DeepPurpose and E3-targetPred methods across different E3-substrate datasets and provided additional benchmarks for our Ubicon model.

### Probability calibration of Ubicon scores

To ensure that Ubicon’s output prediction scores could be reliably interpreted as probabilities, we applied probability calibration techniques. We evaluated two standard calibration methods: isotonic regression^49^ and temperature scaling^50^. temperature scaling adjusts model confidence by dividing the pre-softmax logits by a single learned scalar temperature parameter, while isotonic regression is a non-parametric approach that fits a piecewise constant, non-decreasing function to map raw scores to calibrated probabilities.

The calibration models were trained and evaluated using the prediction scores generated on the held-out test folds during the 5-fold cross-validation procedure used for Ubicon’s primary training and evaluation. Specifically, the prediction scores and corresponding true labels from all 5 test folds were concatenated to create a dedicated calibration dataset. This ensures that the calibrator is fitted on data that the model predicting the scores for that specific fold has not been trained on. This fitting was performed using the isotonic regression implementation available in the scikit-learn Python library (version 1.3.2). For temperature scaling, we implemented the calibration technique optimizing the temperature parameter by minimizing negative log-likelihood on the validation data. The optimization was performed using the BFGS algorithm implemented in SciPy’s optimization library (version 1.10.1). The resulting trained isotonic regression calibrator was saved and subsequently applied to transform all raw Ubicon prediction scores (including those generated for the proteome-wide ESIome atlas in Table S3) into calibrated probabilities for reporting and downstream analyses.

### Generation of the Proteome-wide ESI Prediction Catalog

To generate a comprehensive catalog of potential E3-substrate interactions (ESIs) across the human proteome, we considered all possible pairings between 486 known and putative human E3 ligases and approximately 20,000 reviewed human proteins (sequences ≤ 2046 residues) obtained from UniProtKB^89^.

Recognizing that ubiquitination by an E3 ligase necessitates a physical protein-protein interaction (PPI) with its substrate^96^, we implemented a PPI pre-filtering step prior to applying Ubicon. We employed DeepTrio, a state-of-the-art sequence-based PPI predictor^97^, to enrich the candidate pool for pairs likely capable of direct physical contact.

DeepTrio was applied to predict interaction potential for all possible pairs between the 486 E3 ligases and the ∼20,000 substrate candidate proteins. This PPI pre-filtering significantly reduced the computational burden and focused subsequent high-resolution ESI prediction on pairs with a higher prior probability of biological relevance.

The optimized Ubicon model was then used to predict Ubicon scores. Finally, to ensure interpretability, the raw Ubicon output scores were transformed into calibrated probabilities using the isotonic regression model previously trained on out-of-fold cross-validation data. These calibrated probability scores constitute the final human ESIome prediction atlas provided in Supplementary Table S3.

### Network Analysis

#### Network construction and topological analysis

An E3-substrate interaction network was constructed using human E3 ligases and substrate proteins as nodes. Edges represented predicted ESIs with a calibrated Ubicon score ≥ 0.8; interactions below this threshold were excluded from the topological and community analyses. The resulting graph was treated as undirected. Network construction and graph analyses were performed using the NetworkX library^98^ (version 3.1) in Python. Node degree centrality (number of connections per node) was calculated for all proteins. The top 10 E3 ligases and top 10 substrate proteins ranked by degree were identified as network hubs. The average clustering coefficient of the network was calculated to assess local network density. Network visualization was performed using Cytoscape^99^ (version 3.10.3).

### Community Detection

Densely interconnected subgroups (communities or modules) within the network were identified using the Louvain method for community detection^74^, implemented in Python-Louvain package (community: version 0.16) which optimizes network modularity. Basic statistics, including the number of nodes, E3 ligases, and substrates, were calculated for the major communities detected.

### Functional enrichment analysis

To assess the biological significance of the detected network communities, KEGG pathway enrichment analysis was performed. For the three largest modules identified by the Louvain method, lists of constituent proteins were extracted. Crucially, to avoid biases originating from the functions of the E3 ligases themselves, known E3 ligases were removed from these lists, and only the substrate proteins within each module were used as input for the enrichment analysis. The analysis was conducted using the gprofiler-official Python package^100^ (version 1.0.0) querying the KEGG pathway database^101,102^ for *Homo sapiens*. The background set for enrichment testing consisted of the default g:Profiler organism background. Pathways with an adjusted *p*-value < 0.05, calculated using g:Profiler’s internal g:SCS algorithm for multiple testing correction, were considered significantly enriched. Enrichment results were visualized using −log_10_(adjusted *p*-value), typically showing the top significantly enriched pathways per module.

### Cancer genomics data analysis

TCGA Data acquisition and processing transcriptomic data (RNA-Seq HTSeq counts) and associated clinical information for cancer types relevant to Figure 5 (e.g., Breast Invasive Carcinoma [BRCA], Lung Adenocarcinoma [LUAD]) were obtained from The Cancer Genome Atlas (TCGA) Genomic Data Commons (GDC) portal on March 28, 2025. Data retrieval and preparation were facilitated using the TCGABiolinks R/Bioconductor package^103^ (version 2.30.4). Prior to differential expression analysis, genes with low expression, defined as having a total count of 1 or less across all samples in the respective cohort, were removed to reduce noise.

### Differential gene expression analysis

Differential expression analysis between tumor samples and available normal tissue samples within each TCGA cohort was performed using the DESeq2 R package^104^ (version 1.42.1). DESeq2 models raw counts using a negative binomial distribution and performs normalization based on library size. Log_2_ fold changes (LFC) between tumor and normal conditions were estimated using the apeglm shrinkage estimator^105^ implemented within DESeq2 to stabilize LFC estimates for low-count genes. Genes with a Bonferroni-adjusted *p*-value < 0.05 were considered significantly differentially expressed.

Survival Analysis Kaplan-Meier survival analysis was conducted to evaluate the association between the mRNA expression levels of specific E3 ligases (AURKA, CDC20, UHRF1) and overall survival time in the TCGA BRCA and LUAD patient cohorts. For each gene, patients were stratified into high-expression and low-expression groups based on the median expression value within the cohort. Survival curves were generated and compared using the survival R package^106^ (version 3.5.7) and the log-rank test. A log-rank *p*-value < 0.05 was considered statistically significant.

### Molecular modeling and structural analysis

Initial three-dimensional structures of protein complexes used for structural analysis were predicted using Boltz-1^107^. Structure preparation and refinement were subsequently performed using the Molecular Operating Environment (MOE, version 2020.02; Chemical Computing Group ULC, Montreal, QC, Canada;).

Hydrogen atoms were added to the predicted complex structures using MOE’s standard protonation tools. The structures were then energy minimized using the Amber10:EHT force field^108^ within MOE until a specified convergence criterion was met (RMS gradient < 0.1 kcal/mol/Å^2^). The resulting optimized structures were used for subsequent structural visualization and analysis of interaction interfaces.

### Statistical analysis

General statistical analyses were performed using the scipy.stats module (version 1.10.1) within Python (version 3.8.20). Comparisons of two models across the same cross-validation folds were assessed using a two-sided paired *t*-test. A *p*-value less than 0.05 was considered statistically significant. Statistical methods employed for specific analyses, such as pathway enrichment (e.g., calculation and adjustment of *p*-values by g:Profiler) or survival analysis (e.g., log-rank test), are described in their respective method sections or figure legends.

## ACKNOWLEDGEMENTS

This work was supported by KAKENHI grants from the Japan Society for the Promotion of Science (JSPS) to H.S. (23K28184), as well as Takeda Science Foundation. We thank all the laboratory members for discussion and K. Tanaka for help with preparation of the manuscript.

The results shown here are in part based upon data generated by the TCGA Research Network: https://www.cancer.gov/tcga.

## AUTHOR CONTRIBUTIONS

H.S. conceived of, designed, and supervised the study. T.S and Y.O developed Ubicon and performed all formal analyses. T.S, Y.O. and H.S. jointly wrote the manuscript. All authors have read and approved the final manuscript.

## COMPETING INTERESTS

The authors declare no competing interests.

## Supplementary Information

### Supplementary tables

**Table S1:** Performance evaluation of Ubicon: Stepwise refinement and comparison with existing methods.

| ESI classifier | Features | Mean AUROC ± SD | Mean AUPRC ± SD | Mean F1 Score ± SD | Ref |
| --- | --- | --- | --- | --- | --- |
| E3-targetPred | Sequence only | 0.6745 ± 0.0185 | 0.1785 ± 0.0160 | 0.2311 ± 0.0092 | 29 |
| DeepUSI<br>(DeepPurpose) | Sequence only | 0.8485 ± 0.0049 | 0.4445 ± 0.0174 | 0.3944 ± 0.0294 | 28, 94 |
| CNN-based | ESM-C 6B + 3Di | 0.8262 ± 0.0080 | 0.3143 ± 0.0316 | 0.3405 ± 0.038 | This study |
| Catboost | ESM-C 6B + 3Di | 0.9136 ± 0.0040 | 0.5901 ± 0.0185 | 0.4480 ± 0.0212 | This study |
| Catboost | ESM-C 300M + 3Di | 0.9135 ± 0.0044 | 0.5939 ± 0.0139 | 0.4744 ± 0.0253 | This study |
| Catboost | LoRA-tuned ESM-C 300M + 3Di | 0.9291 ± 0.0063 | 0.6771 ± 0.0169 | 0.6073 ± 0.0198 | This study |
| Catboost | LoRA-tuned ESM-C 300M + Subcellular Localization | 0.9287 ± 0.0062 | 0.6737 ± 0.0132 | 0.5816 ± 0.0127 | This study |
| Catboost<br>(Ubicon) | LoRA-tuned ESM-C 300M + 3Di + Subcellular Localization | 0.9305 ± 0.0035 | 0.6812 ± 0.0161 | 0.6210 ± 0.0158 | This study |

**Table S2:** Ubicon prediction scores for known orthologous E3-substrate interactions in *Mus musculus*.

| E3 ligase | Substrate | UniProtID (E3) | UniProtID (Substrate) | Ubicon score |
| --- | --- | --- | --- | --- |
| β-TrCP | p53 | Q5SRY7 | P02340 | 0.88333 |
| Rnf4 | Parp1 | Q9QZS2 | P11103 | 0.87805 |
| Fbxo21 | Ccnb1 | Q8VDH1 | Q3TQW9 | 0.82171 |
| Fbxw7 | Myc | Q8VBV4 | P01108 | 0.82170 |
| Mdm2 | P53 | P23804 | P02340 | 0.72727 |
| Rnf5 | STING1 | O35445 | Q3TBT3 | 0.68639 |

**Table S3: Comprehensive Ubicon prediction scores for the human E3-substrate interactome (ESIome). (provided in CSV format)**

**Table S4:**
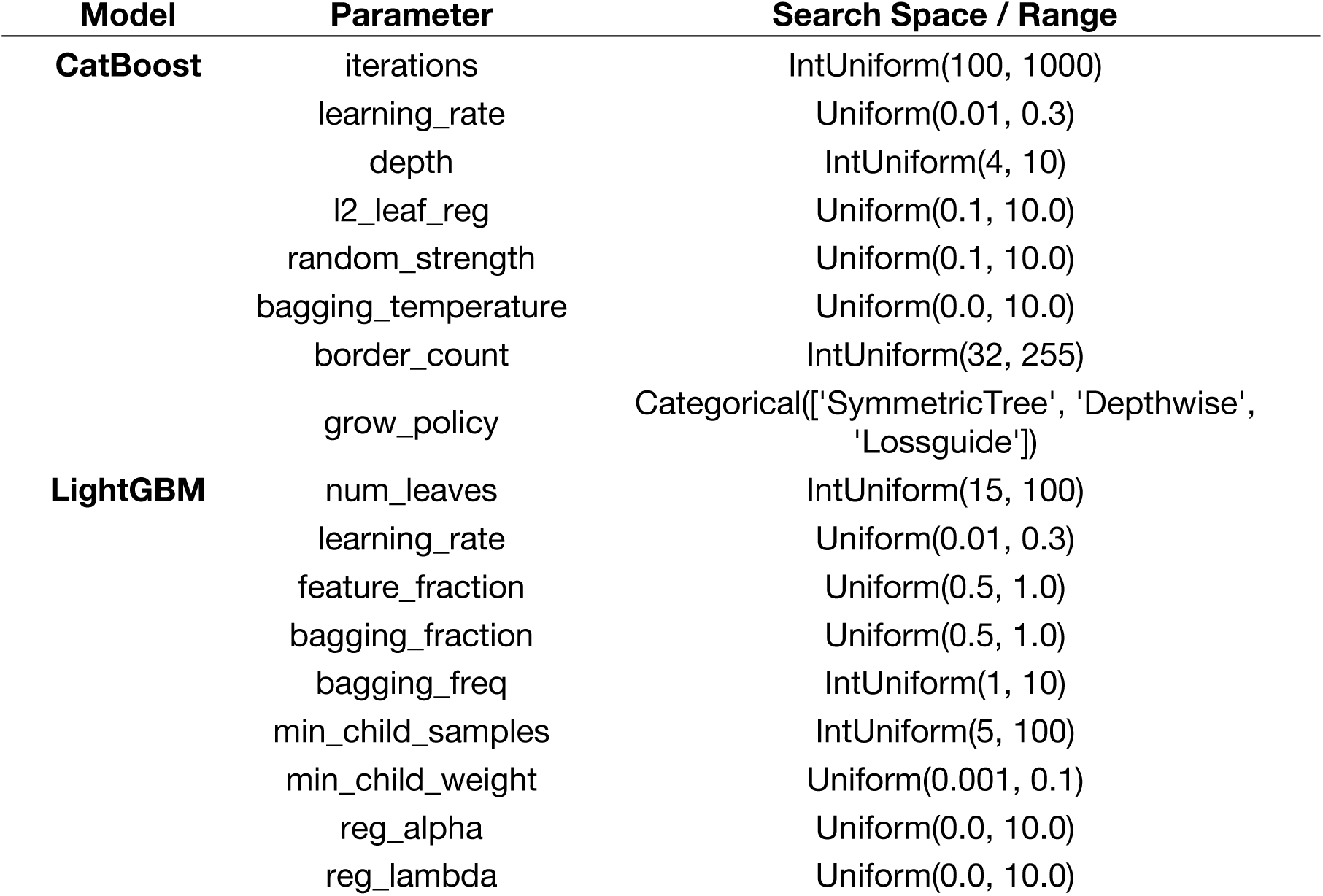
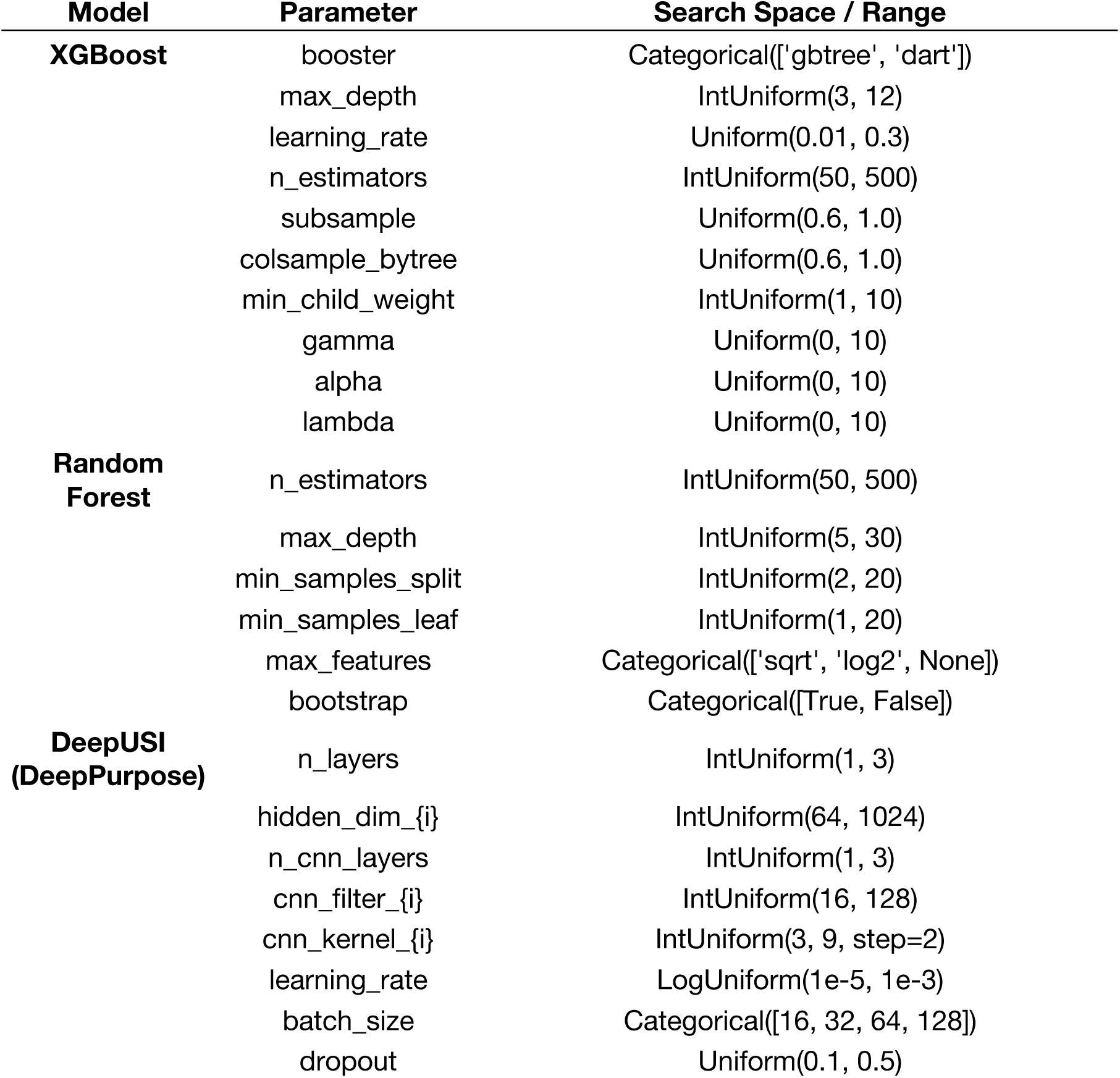
Optuna hyperparameter search spaces for ESI prediction models.

### Supplementary figure legends

**Figure S1.**
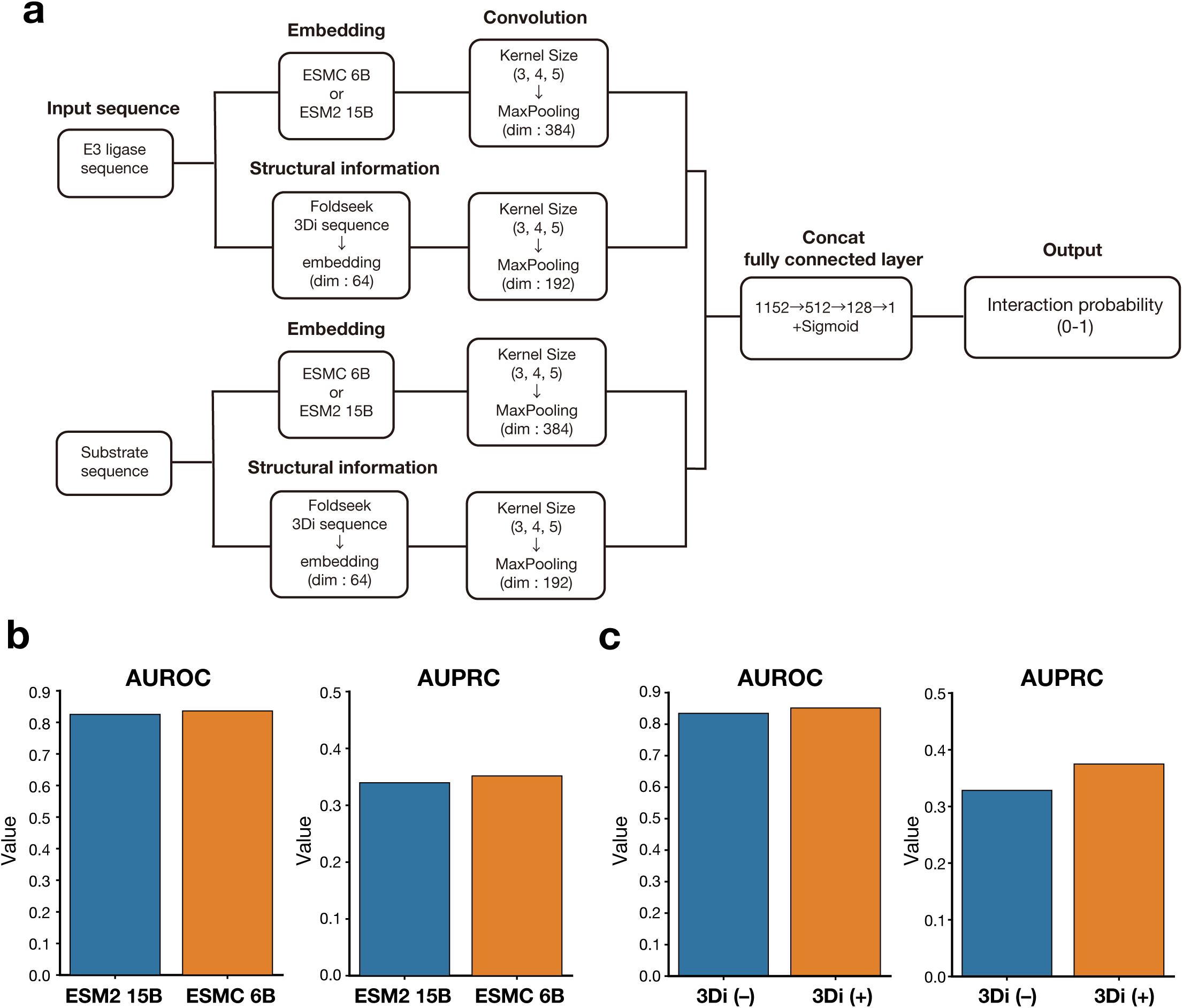
Initial model architecture, feature selection, and evaluation for Ubicon development. (**a**) Schematic of the initial CNN-based architecture evaluated for Ubicon. Inputs are E3 ligase and substrate amino acid sequences. Features include sequence embeddings derived from protein language models (ESM-C 6B or ESM2 15B) and structural information represented as 3Di sequences generated from AlphaFold2 structures via Foldseek. Sequence and structural features are processed through separate convolutional and pooling layers before concatenation and final prediction of the E3-substrate interaction probability by fully connected layers. (**b**) Performance comparison between protein language models ESM2 (15B parameters) and ESM-C (6B parameters) for ESI prediction, using sequence information only (no 3Di features). Bar plots show the Area Under the Receiver Operating Characteristic Curve (AUROC) and Area Under the Precision-Recall Curve (AUPRC). ESM-C (6B) was selected over ESM2 (15B) based on comparable predictive performance at lower computational cost and its capacity to handle longer protein sequences, enabling wider proteomic coverage. (**c**) Evaluation of the contribution of structural features to prediction performance. Bar plots compare AUROC and AUPRC for the ESM-C (6B) model using sequence embeddings alone (3Di (-)) versus using both sequence embeddings and 3Di structural information (3Di (+)). The inclusion of 3Di structural features derived from AlphaFold2/Foldseek improved model performance, particularly AUPRC.

**Figure S2.**
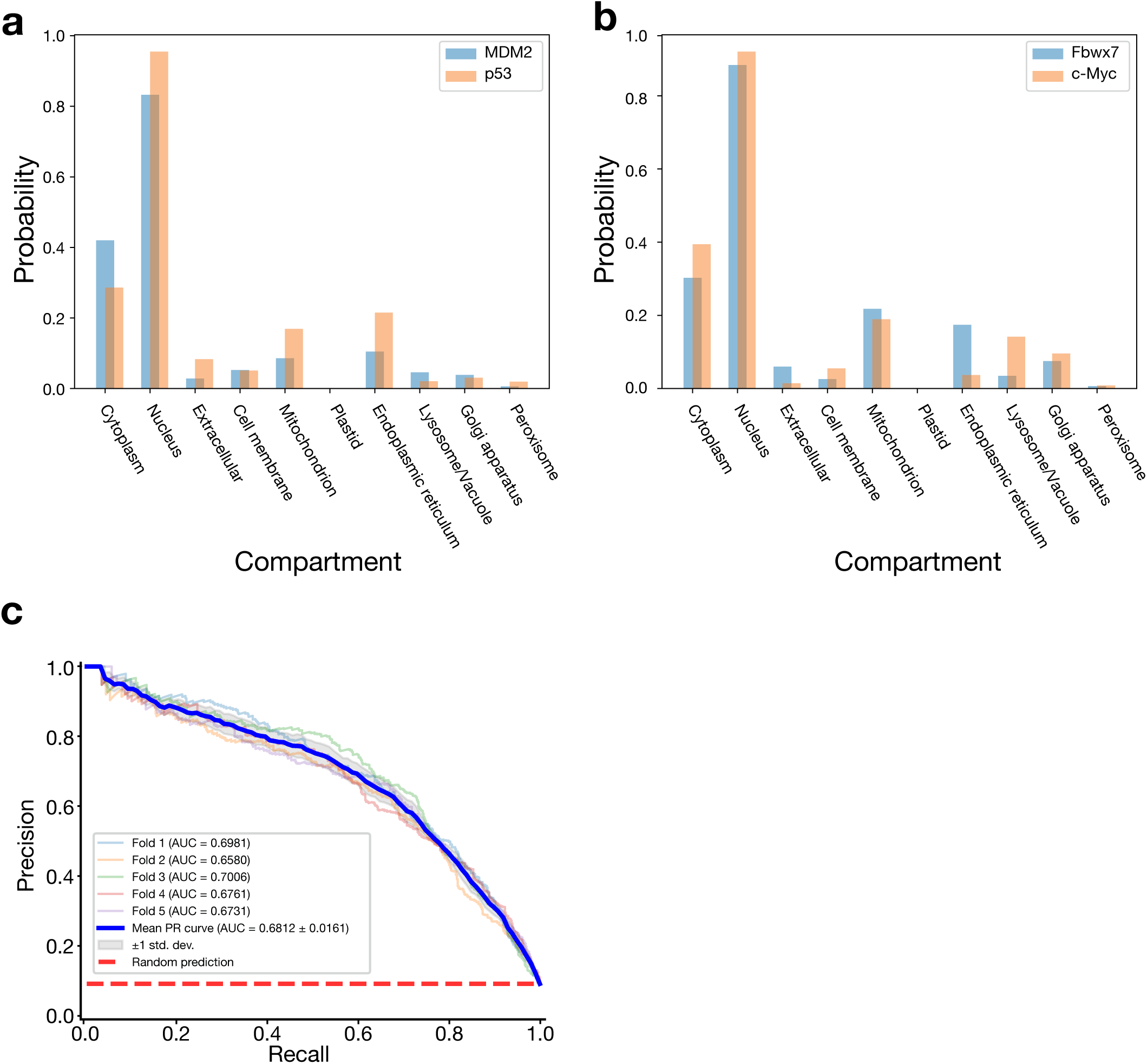
Validation of localization features and Precision-Recall performance of the Ubicon model. (**a**, **b**) Validation of subcellular co-localization for known ESI pairs. Bar plots show predicted localization probabilities (obtained using DeepLoc2) across 10 cellular compartments for the components of two known interacting pairs: (a) MDM2 and p53, and (b) Fbxw7 and c-Myc. High predicted probabilities of co-localization in the Nucleus and/or Cytoplasm for both pairs support the biological rationale for using localization information as predictive features in Ubicon. (**c**) Precision-Recall (PR) curves from 5-fold cross-validation of the final Ubicon model. Individual fold PR curves, the mean PR curve (bold blue line), and the baseline performance expected by random prediction (dashed red line) are shown. The high mean Area Under the PR Curve (AUPRC = 0.6812 ± 0.0161) demonstrates Ubicon’s ability to reliably identify true positive interactions with high precision, which is particularly important for discovery applications on imbalanced datasets typical for ESI prediction.

**Figure S3.**
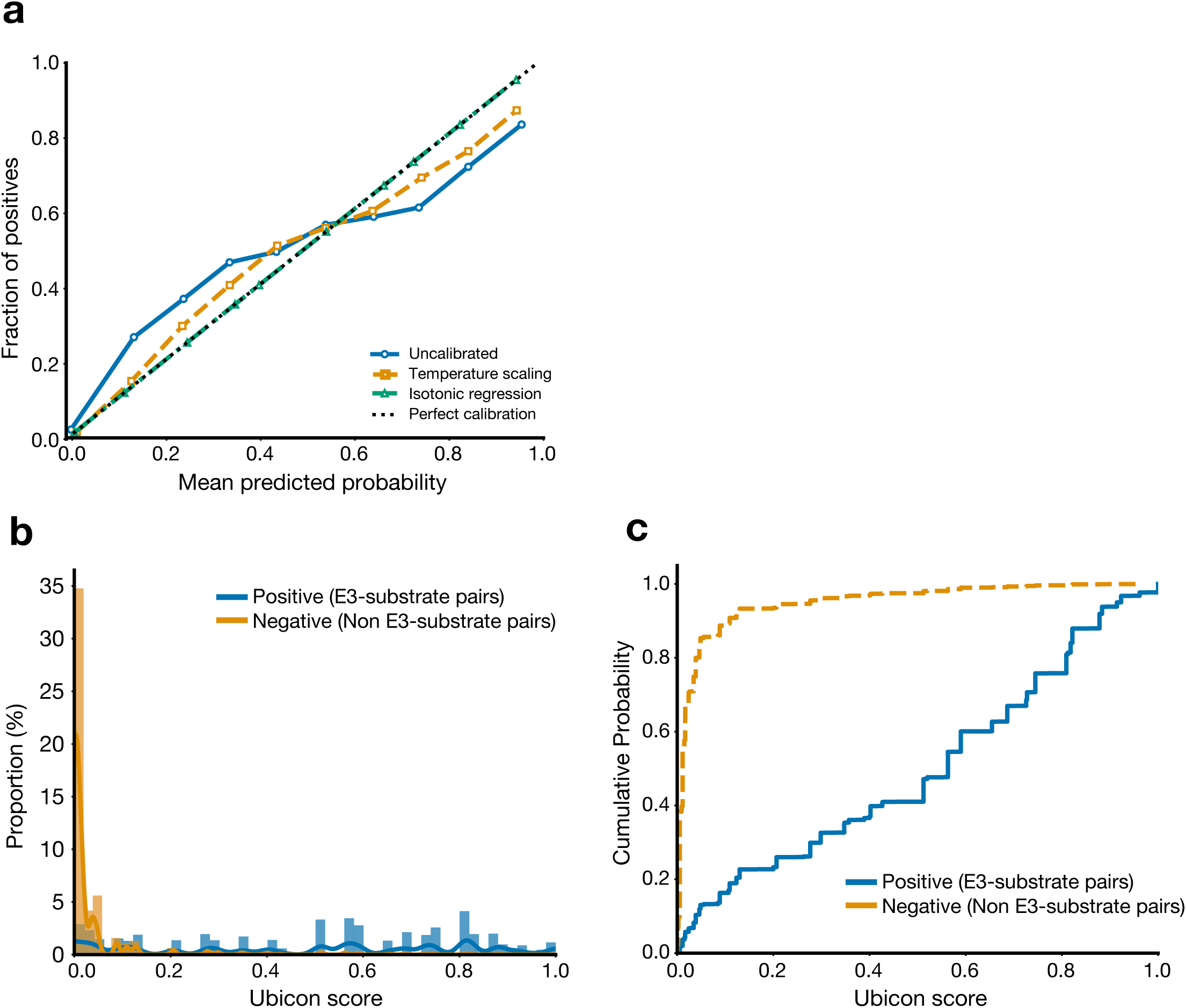
Improving interpretability of Ubicon prediction scores through probability calibration. (**a**) Reliability diagram evaluating probability calibration methods for Ubicon scores by comparing the relationship between predicted probability bins (x-axis) and the observed fraction of positive interactions within each bin (y-axis) for uncalibrated scores (blue), temperature scaling (orange), and isotonic regression (green) against perfect calibration (dashed diagonal). isotonic regression yields the best-calibrated probabilities, closely matching the ideal diagonal line. (**b**) Distribution of Ubicon scores after calibration with isotonic regression (Calibrated Score (IR)). Histograms show distinct distributions for positive (blue, E3-substrate pairs) and negative (orange, Non E3-substrate pairs) interactions, with positive cases enriched at higher scores. (**c**) Cumulative distribution functions (CDFs) for isotonic regression-calibrated Ubicon scores, shown separately for positive (blue) and negative (orange) interactions. The separation between the CDFs highlights the discriminative power of the calibrated scores.

**Figure S4.**
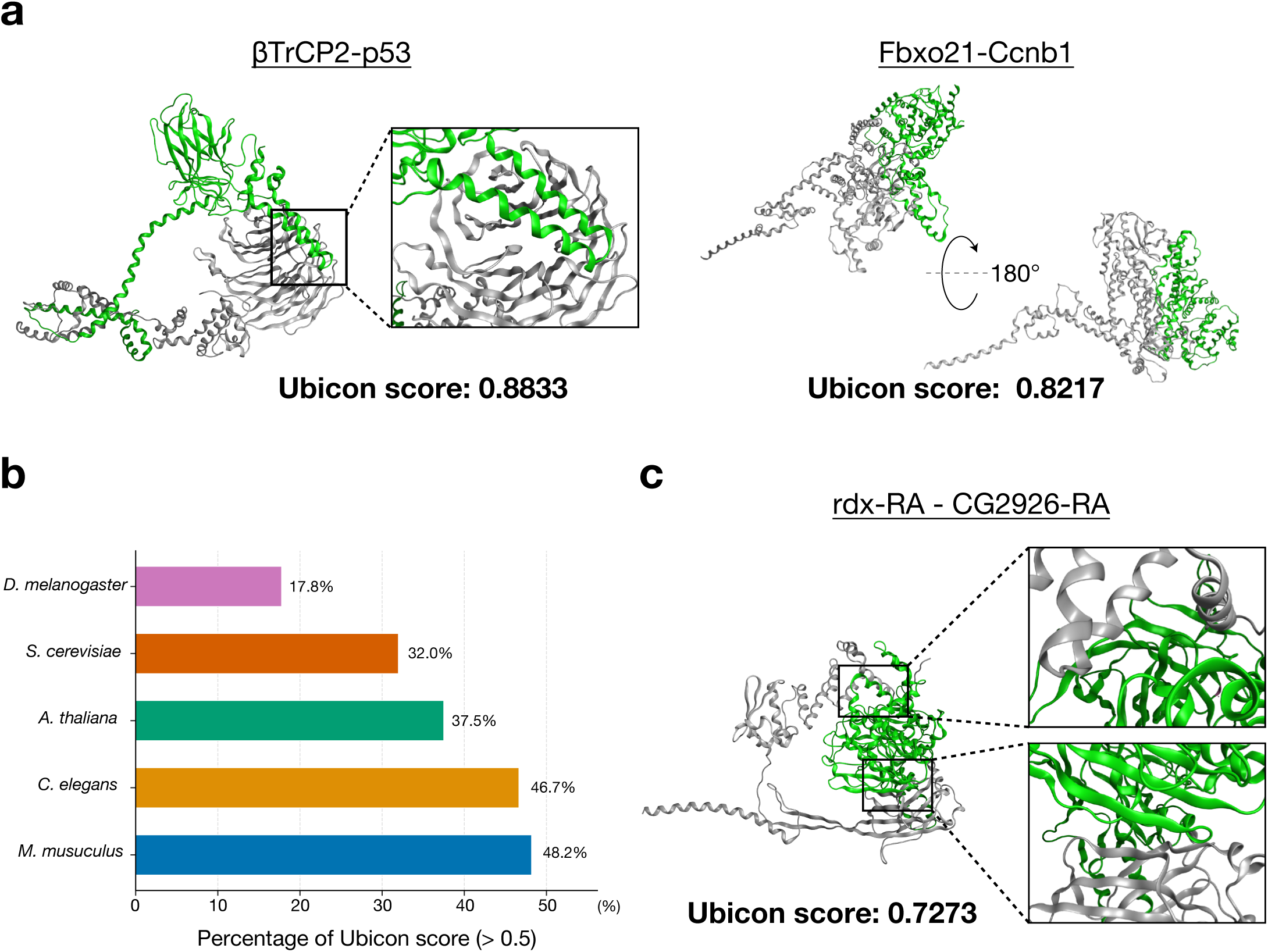
Ubicon captures conserved ESI recognition principles across eukaryotes. (**a)** Ubicon accurately predicts known orthologous ESIs in *Mus musculus*. Predicted complex structures (E3 ligase: gray; substrate: green) and high Ubicon scores are shown for the βTrCP2-p53 (score: 0.8833) and Fbxo21-Ccnb1 (score: 0.8217) interactions. (**b)** Ubicon performance on curated ESI datasets across diverse eukaryotes. Bar chart shows the percentage of known ESI pairs assigned a calibrated Ubicon score > 0.5 for *M. musculus*, *D. melanogaster*, *C. elegans*, *S. cerevisiae*, and *A. thaliana*, indicating broad generalization capacity. (**c)** Example of cross-species prediction for the *Drosophila melanogaster* ESI pair rdx-CG2926 (homologous to human SPOP-CCNB1). The predicted complex structure (E3 ligase: gray; substrate: green) and the assigned Ubicon score (0.7273) are shown.

## Notes

### Competing Interest Statement

The authors have declared no competing interest.

